# Bioorthogonal antigens allow the unbiased study of antigen processing and presentation

**DOI:** 10.1101/439323

**Authors:** Can Araman, Linda Pieper-Pournara, Tyrza van Leeuwen, Arieke S. B. Kampstra, Thomas Bakkum, Mikkel H. S. Marqvorsen, Clarissa R. Nascimento, G. J. Mirjam Groenewold, Willemijn van der Wulp, Marcel G. M. Camps, Herman S. Overkleeft, Ferry A. Ossendorp, René E. M. Toes, Sander I. van Kasteren

## Abstract

Proteolysis is fundamental to many biological processes. In the immune system, it underpins the activation of the adaptive immune response: degradation of antigenic material into short peptides and presentation thereof on major histocompatibility complexes, leads to activation of T-cells. This initiates the adaptive immune response against many pathogens.

---

Studying proteolysis is difficult, as the oft-used polypeptide reporters are susceptible to proteolytic sequestration themselves. Here we present a new approach that allows the imaging of antigen proteolysis throughout the processing pathway in an unbiased manner. By incorporating biorthogonal functionalities into the protein in place of methionines, antigens can be followed during degradation, whilst leaving reactive sidechains open to template and non-templated post-translational modifications, such as citrullination and carbamylation.

Using this approach, we followed and imaged the in cell fate of the commonly used antigen ovalbumin, as well as the post-translationally citrullinated and/or carbmhyalted auto-antigen vinculin in rheumatoid arthritis.

Proteolysis underpins many fundamental biological processes at every developmental stage: from the essential role of the protease fertilin during fertilization^1^ to the initiation of cell death through the activation of caspases^2^. It is therefore unsurprising that dysregulation of proteolysis is a hallmark in a variety of pathologies: aberrant protease activity has been implicated in the growth and metastasis of tumors^3^, auto-immune pathogenesis^4–5^, compromised bone desorption^6^, and intracellular bacterial persistence^7^, to name but a few.

The biology of proteases is complex: factors such as their compartmentalized distribution^8^, post-translational activation^9^, functional redundancy and interplay^10^ complicates their study. This is none the more prominent than during antigen processing by antigen presenting cells (APCs)^11–12^. APCs take up exogenous material, degrade the proteins from these sources and present specific peptides from the degraded protein on their major histocompatibility complexes (MHCs)^13^. Recognition of a specific peptide-MHC (pMHC) by a cognate T-cell then leads to the initiation of the adaptive immune response against the source of the peptide^13^. This process is of prime importance in the clearance of exogenous pathogens and cancer, but also for the opposite: the induction of tolerance against innocuous substances and self-tissue^14–16^. As such, the understanding of the proteolysis underpinning this phenomenon is essential.

Proteolysis in APCs is complex: multiple protease and peptidase families (e.g. aspartic, serine, and cysteine proteases) are involved in the production of MHC-restricted peptides^17–19^. Many of these proteases, particularly those in the endo-lysosomal system, are under tight post-translational control^9^: they are produced as zymogens, routed to the vesicles of the endolysosomal compartment where they are activated by a combination of the low pH and the removal of inhibitory peptides through the activity of other proteases^9^. The often-promiscuous substrate preference by virtue of their shallow binding grooves^20^, attenuation of their activity by endogenous inhibitors^21^, changes in pH, and radical concentrations^22^ are factors that further complicate the study of these enzymes.

The nature of the antigen (i.e. the substrate) itself also influences proteolysis^23^ by altering protease specificity or overall stability. For example, a point mutation in the multiple sclerosis (MS) auto-antigen myelin basic protein (MBP_85-99_) prevents its cleavage by the non-papain-like cysteine protease asparagine endopeptidase, leading to enhanced presentation to T-cells^4^. (De)stabilizing a protein fold can also alter antigenicity: the sequentially identical, but structurally destabilized variants RNAse S and apo-HRP of the stable proteins RNAse A and HRP are both poor antigens compared to the stable counterparts. They are rapidly degraded *in vitro,* and antigen presentation efficiency is significantly reduced *in vivo*^24^. Stabilizing a fold, on the other hand can also reduce its antigenicity, as observed for hen egg lysozyme^25–26^.

Post-translational modification of the antigen too can alter proteolysis by altering specificity. This can, for example, lead to the appearance of ‘neo-auto-antigens’ – non-thymic peptides that can drive auto-immune diseases – and has been observed in MS: an arginine to citrulline modification in myelin oligodendrocyte glycoprotein in B-cells prevented destructive epitope processing of such a neo-auto-antigen, leading to T-cell activation in a marmoset model of the disease^5^.

The go-to approach for studying antigen processing has been the use of model antigens and looking at the downstream presentation of immunodominant peptides from these antigens. This method does not yield information on the subcellular events that govern the processes underpinning this T-cell activation. Alternatively, detectable model proteins have been used to study subcellular routing, such as lactamases^27^, luciferase^28^, and HRP^29^. These proteins have proven valuable in the study of early events in the process using enzymatic activation as a biochemical read-out. However, the essential role for proteolysis during antigen presentation – which will render these reporters catalytically inactive, precludes their use for obtaining information from later stage events.

For the study of these late stage processing events, covalent fluorophore-antigens adducts have been used^30–32^. The low molecular weight (∼ 800 Da), and stability to proteolysis of the fluorophores are beneficial traits of these reagents. However, the change in charge, hydrophobicity/aromaticity, shape and structure can alter the properties of the protein^33–34^ which we hypothesize will lead to alterations in processing and antigenicity.

We here explore bioorthogonal chemistry^35^ as an approach to study antigen degradation in immune cells circumventing the aforementioned downsides (Fig. 1A). Bioorthogonal chemistry is the performance of a selective chemical ligation reaction in the context of the biological milieu^36–38^. A small abiotic group, such as an azide or alkyne is first introduced into a biomolecule and can then – when the biological time course is completed – be selectively modified with a detectable group (e.g. fluorophores or biotin) for identification. We envisioned that a bioorthogonal antigen – in which the antigen contained these modifications - would be highly suitable for studying degradation events during antigen presentation: the labels introduced into proteins are minimally altered compared to naturally occurring amino acids, and bioorthogonal groups are available that are fully stable to the conditions found during endo/phagosomal maturation^39^. Most importantly, recombinant proteins can readily be obtained in good yields that allow the execution of functional immune assays that contain multiple copies of the ligation handles azidohomoalanine and homopropargylglycine (Aha/Hpg, Fig. 1) in place of Met-residues^40–42^, without loss of their activity^43^.

**Figure 1:**
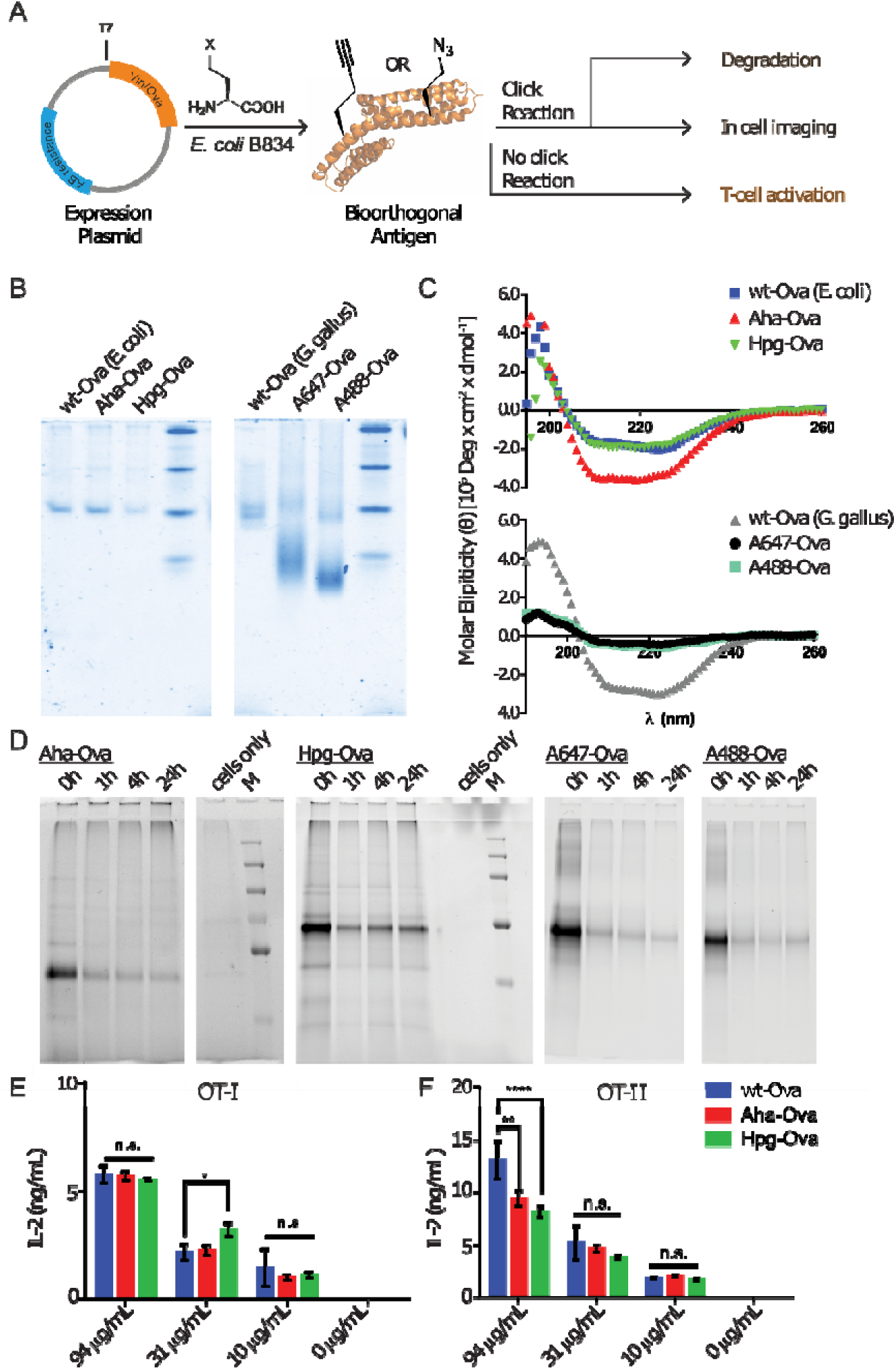
Bioorthogonal antigens closely approximate the biophysical behavior of wild type antigens. A) Expression of antigenic proteins in the *E. coli* auxotrophic strain B834 allows replacement of methionines with azidohomoalanine (Aha) or homopropargylglycine (Hpg). These labels allow the in cell imaging of degradation whilst at the same time allowing the study of the antigens by T-cell activation; B) native-PAGE analysis shows similar migration behavior of wt-Ova, Aha-Ova and Hpg-Ova, whilst showing altered migration of fluorophore modified Ova-analogues; C) circular dichroism of bioorthogonal and fluorophore Ova-analogues; D) tracking of degradation of antigen inside D1 dendritic cells; E) activation of OT-I cells by Ova-variants as measured by IL-2 production; F) activation of OT-II cells by these same antigens. Asterisks refer to given P values, * p < 0.05, ** p < 0.005, **** p < 0.00005. Group mean values were analyzed by two-way analysis of variance with Tukey post hoc significant difference test using GraphPad Prism 6.0. Data are represented as means ± SD.

In this study we produce, characterize and explore bioorthogonal antigens to follow degradation inside the APC whilst only minimally impacting on their secondary structure and the rate/nature of degradation. Furthermore, we want to use these reagents to study the effect of pathogenic PTMs such as citrullination and carbamylation on proteolysis of an autoimmune disease (RA) related antigen, vinculin (Vin).

## Results

We produced methionine-containing and bioorthogonally labelled variants of the model antigen ovalbumin (Ova) in the methionine auxotrophic strain B834 of *E. coli*^44^. This strain allows for the facile incorporation of unnatural amino acids in place of methionine residues by placing the bacterial strain in a methionine-free expression medium augmented with Aha or Hpg^40,42^. Ova-containing Aha amino acids (Aha-Ova) at all 17 naturally occurring Met-codon positions was produced from the plasmid pMCSG7 that contains the Ova-gene fused to a Hexa-His-Tag (His_6_), *via* a TEV cleavage site (a kind gift from N. del Cid and M. Raghavan^45^) using existing protocols^46–47^. Ova containing homopropargylglycine (Hpg-Ova) or methionine (wt-Ova) at these positions were also produced. The yields of expression were 2-3 mg/L culture after purification for both wild type and bioorthogonal variants. LC-MS analysis of the bioorthogonal variants indicated quantitative incorporation of the bioorthogonal handles during expression (Fig. S1).

For comparison, we purchased Ova from chicken egg (wt-Ova *G. gallus*) – which is different from our expressed Ova in its absence of a His-tag and the presence of one (or in rare cases two) glycans, mainly of high mannose/hybrid glycoforms^48^ – as well as fluorophore-modified variants of this chicken egg Ova (A647-Ova and A488-Ova).

We compared the various bioorthogonal and fluorophore modifications in terms of structure and stability. Native PAGE analysis (Fig. 1B) showed equal gel migration properties for bacterially expressed and isolated Ova analogues. The bioorthogonal variants also showed identical migration profiles. Fluorophore modified A647-Ova and A488-Ova on the other hand, showed several bands and distinct migration profiles compared to purchased and bacterially expressed Ova variants. Circular dichroism (Fig. 1C, CD) of Ova variants confirmed these structural similarities and differences: a drop in elipticity was observed for A647-and A488-Ova (Fig. 1C, lower graph), indicating a lower structural stability. Hpg-Ova showed identical elipticity and Aha-Ova even an enhanced elipticity compared to wild type, indicating an increase in structural stability.

To assess whether the bioorthogonal variants were processed similarly to wild type, the *E. coli* expressed Ova-variants were subjected to proteolysis by isolated lysosomal extracts from the macrophage cell line RAW264.7, which is highly proteolytically active^49^. No processing of Ova was observed under these conditions, even after prolonged (24 h) incubation (Fig. S2), something that was also observed previously for various dendritic cells, but not for primary macrophages^50^. These data suggest that, for ovalbumin, digestion by isolated lysosomes is not a suitable model for the degradation.

We thus proceeded to study whether the bioorthogonal Ova-variants could be used to show the degradation in live APCs (Fig. 1D). As the antigen and its breakdown fragments have to be selectively detected within the APC-lysates, only fluorophore or bioorthogonal antigens can be used for this purpose. To test the degradation of these antigens, D1 APCs^51^ were pulsed with antigen for 2 h (Aha-, Hpg-Ova, A488-Ova and A647-Ova). The cells were then washed and chased for 0 h, 1 h, 4 h or 24 h. After fixation, the cells were lysed and fluorophores introduced by a copper-catalyzed Huisgen cycloaddition (ccHc)-reaction with A647 azide/alkyne. Labelled lysates were then separated by SDS-PAGE and the fluorescent signal was visualized after gel electrophoresis. In contrast to the stability in isolated lysosomes, degradation for these variants in D1s did occur. For all samples a decrease in the band for intact Ova was observed, but no intermediate-sized fragments. The small peptide/amino-acid-sized fragments cannot be visualized for the bioorthogonal antigens, as the excess of unreacted fluorophore obscured all bands < 3 kDa (which was thus removed from these gels). In theory, A647-Ova should allow the visualization of low molecular weight species, as this protein was purified to remove unreacted fluorophore prior to the biological experiment. However, none such bands are observed for this antigen. This suggests the terminal degradation-fluorophore fragments diffuse away, or that the fluorophore itself does not survive the conditions in the lysosome.

To assess the suitability of bioorthogonal amino acids to study antigen presentation, we also assessed their ability to activate OT-I (CD8^+^) and OT-II (CD4^+^) (Fig. 1E/F) T-cells in co-culture with murine BMDCs. BMDCs were incubated for 4 h with antigens, excess antigen was washed away before co-culture with OT-I or OT-II T-lymphocytes isolated from mouse spleen. After 24 h, activation of T-cells was measured by IL-2 ELISA^52–53^. This assay showed that Aha- and Hpg-Ova variants show similar CD8 T-cell activation compared to *E. coli*- expressed wt-Ova (Fig. 1E). CD4 T-cell activation differed significantly only at the highest Ova-concentration; with the difference in presentation dissipating at the lower concentrations (Fig. 1F). Fluorophore labeled antigens A647-and A488-Ova were also used to activate OT-I and OT-II T-cells, however the experimental variation between replicates was extensive, with the difference between Ova and the fluorophore modified variants varying over two orders of magnitude between repetitions for both OT-1 and OT-II activation assays (Examples shown in Fig. S3).

We also imaged the fate of one of the bioorthogonal antigens, Aha-Ova, as well as A647-Ova by confocal microscopy (Fig. 2, Fig. S4-5 and Supplementary movie 1). For this we pulsed the dendritic cell lines DC2.4^54^ (Fig. 2) and the aforementioned D1s^51^ (Fig. S4) cell lines with the two Ova-variants or with 10 mM NaHCO_3_ buffer as a negative control (< 5 % v/v, Fig. S5, Supporting Information) for 2 h in presence of LPS to allow for maturation of the endosome to a late endosome/lysosome. Surprisingly, only A647-Ova localized partly to (late) LAMP-1 positive vesicles at this time point, whereas Aha-Ova showed no overlap with LAMP-1. Increased uptake of fluorphore-modified antigen was also observed in both the DC2.4 and D1-cell lines. This suggests a possible interference of the fluorophore handle on the routing of the antigen (Fig. 2B), perhaps due to the charge and lipophilicity-induced destabilization of the antigen structure for A647-Ova.

**Figure 2:**
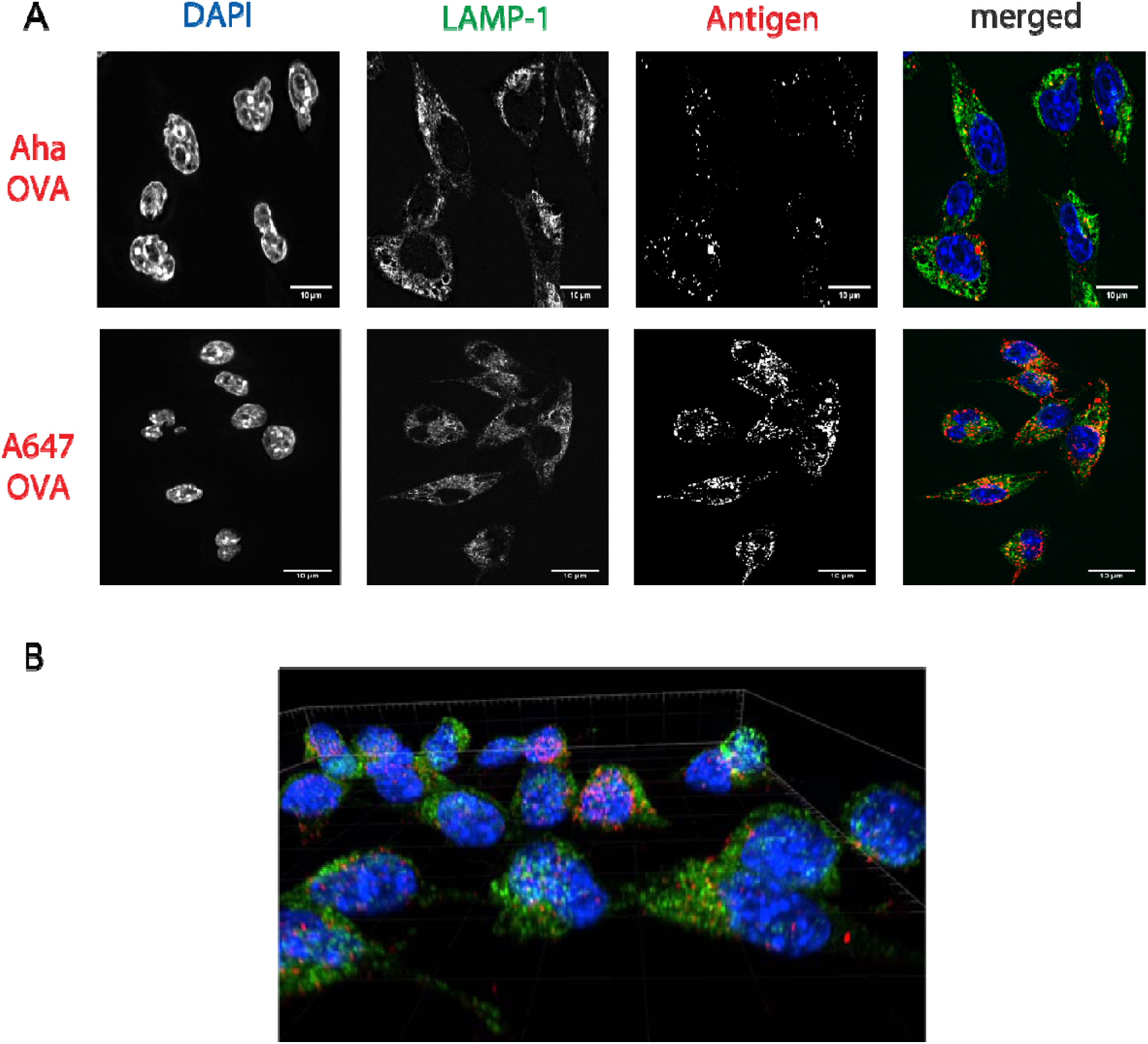
Imaging processing of bioorthogonal and fluorophore-modified Ova in DC 2.4-APCs via confocal microscopy. DCs were incubated for 2 h (pulse) with Aha or A647-Ova. Cells were fixed with 0.5% PFA and processed for immunofluorescence with LAMP-1 as a lysosomal marker (green in merged images). The nucleus was stained with DAPI (blue in merged images). Aha-Ova was stained using ccHc with A647-alkyne fluorophore (grey in single channel, red in merged images). A) upper panel: imaging of Aha-Ova, lower panel: imaging of A647-Ova B) Screenshot from Supplementary Movie 1. Scale bar is 10 µm (white bar, right corner).

These data suggest that bioorthogonal antigens are suitable reagents for the study of antigen processing and presentation, due to the similar structure and biophysical behavior compared to wild type, combined with the ability to introduce a detectable group at the end of the biological experiment.

To apply the technique to a disease relevant system, we turned our focus to the role of lysine post-translational modification on antigen presentation. In rheumatoid arthritis, lysine carbamylation and arginine citrullination have been strongly implicated in the pathogenesis of the disease. In over 60% of patients suffering this disease, mature antibodies against citrullinated^55–57^ and carbamylated proteins are found^58–60^, which suggests that T-cell help to the auto-antibody producing B-cell has taken place. This in turn suggests that carbamylation and citrullination alters the processing and presentation of CD4 restricted neo-epitopes.

To study this phenomenon with bioorthogonal antigens, we expressed a truncated variant of Vinculin (Vin) comprising aa 453-724 of the native sequence containing the potential immune-dominant epitope DERAA^61–62^. This protein was fused to an N-terminal Deca-His-Tag (His_10_) *via* an Enterokinase cleavage site (DDDDDKH). Vin contains 8 internal Met-codons. Bioorthogonal expression yielded a mixture containing mainly 7 or 8 copies of Hpg or Aha. Minor variants with fewer (6, 5 and 4) copies of Aha and Hpg were also detected (combined abundance <10%; Fig.S6).

Bioorthogonal antigens carry their label in place of Met-residues leaving the nucleophilic lysine sidechain free for post-translational modification. We therefore produced carbamylated and/or citrullinated variants of the proteins using methods described previously^63–64^. All Vin variants used in this work contained 23 possible carbamylation sites (incl. the N-terminus). Results of SDS-PAGE and LC-MS measurements showed that >19 amines were carbamylated for all variants of Vin (Fig. S7A/B) as determined by LC-MS. *In vitro* citrullination using recombinant human PAD4, an enzyme that is responsible for citrullination *in vivo*, yielded quantitative Arg ➔ Cit conversion (Fig. S7A/B) for all wt and bioorthogonal variants of Vin.

Carbamylation of the Vinculin variants kept the structure identical to wild type, and the rigidity of the structure (as determined by CD) was identical to the wild type. A small increase in structural stability for Aha-Vin was observed and no change in stability upon carbamylation of Hpg-Vin (Fig. S7C). Citrullination of the Vin-derivatives had a different effect on the structure: these proteins became high in β-sheet content and random coil-rich structure with a decrease in the content of α-helices (Fig. S7C).

We next studied the influence of the structural effect of carbamylation and citrullination on the *in vitro* proteolytic stability degradation of the bioorthogonal antigens. The PTM-modified Vin variants were subjected to *in vitro* lysosomal degradation as described above. In this assay, carbamylated wt-, Aha- and Hpg-Vin-variants all showed a greater resistance to proteolysis than their non-modified counterparts (Fig. 3). At t = 18 h, all non-carbamylated Vin variants had been degraded >50%, as determined by densitometry, whereas the carbamylated Vin-variants were still intact at this time point, and even after 48 h (<10% degradation observed and <30% for carbamylated wt-Vin). Citrullination, despite the induced changes in structure, had no significant effects on the rate of degradation (Fig. S8).

**Figure 3:**
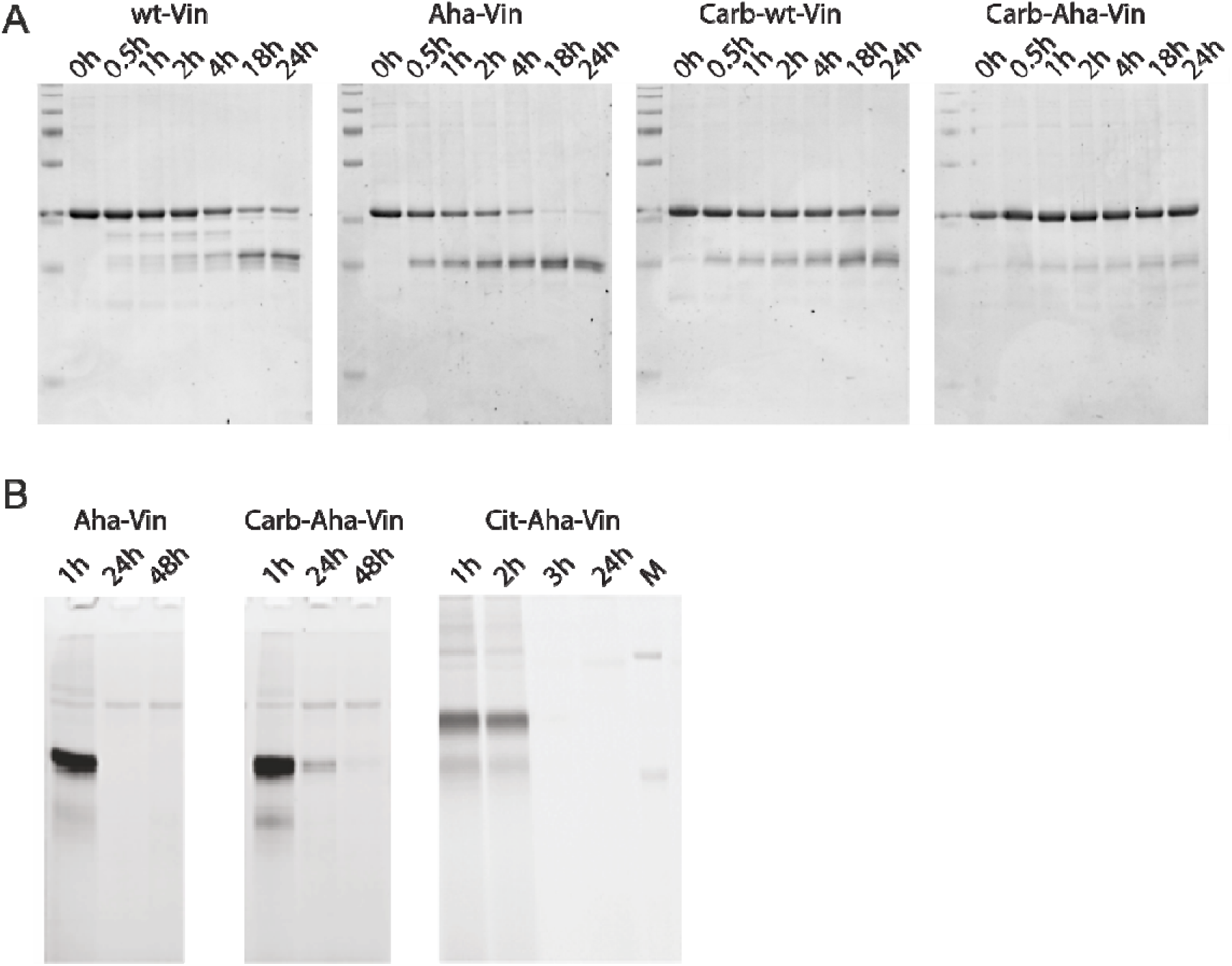
Effect of post-translational modifications on vinculin degradation. A) incubation of vinculin variants with isolated lysosomes show decreased proteolysis of wt and Aha-Vin upon carbamylation; B) imaging this degradation inside dendritic cells (BMDCs) further confirms this stabilizing effect: At t = 24 h only carbamylated Aha-Vin remains visible.

We again determined whether this increased resistance to proteolysis *in vitro* translated to an increased stability in cells. DCs were pulsed with the PTM Vin-antigens for 2 h and then chased for 1 h, 24 h or 48 h respectively. After harvest, lysis, and a ccHc reaction to introduce the appropriate fluorophore, the labelled lysates were separated by SDS-PAGE and the signal stemming from the bioorthogonal groups measured using in-gel fluorescence (Fig. 3B and Fig. S8D/E). Wild type variants of the protein were not visible due to the absence of ccHc-reactive groups (Fig. S9). At t = 24 h none of the antigens were visible, suggesting complete degradation at this time point. The same was observed for citrullinated variants (Fig. 3B). Carbamylated antigens on the other hand showed a clear signal at t = 24 h for all antigens, with a detectable signal even observed at t = 48 h for carbamylated Hpg-Vin (Fig. 3 and Fig. S8D), showing a clear persistence of the protein upon carbamylation in live dendritic cells.

In a final test, whether the approach could also be used for T-cell activation, we tested whether the PTM-modified variants of Vin altered T-cell activation. As a vin-specific T-cell we used the only available T-cell reagents available against a vinculin-antigen, a Jurkat T-cell line (J76) transduced with a T-cell receptor (TCR) restricted to a specific peptide present in vinculin (REEVFDERAANFENH; VCL-DERAA)^65^. The resultant reporter cells allow simultaneous determination of the activity of the transcription factors NF-κB, NFAT and AP-1 that play key roles in T-cell activation. The percentage of fluorescence positive cells after antigen-specific activation can be readily measured using flow cytometry.

The Jurkat-DERAA cells were first stimulated by a HLA-DQA1*03:01/DQB1*03:02+ (HLA-DQ8+) EBV-transformed B cell line loaded with titrated amounts of the vinculin peptide recognized by the TCR. The Jurkat T-cells showed a peptide dose-dependent response as determined by eCFP expression as read-out (Fig. 4). The response was HLA-DQ8-restricted as HLA-DQ8-negative B cells were unable to stimulate the T-cells (Fig. S10). Next the monocyte-derived dendritic cells, isolated from HLA-DQ8+ donors, were pulsed o.n. with different bioorthogonal proteins and their modified counterparts (3 µM). After DC maturation for 30 h, the Jurkat T-cells were added and used as read-out for HLA-restricted T-cell activation by flow cytometry (Fig. 4 and Fig. S10-11). Carbamylation also did not affect antigen presentation, despite its strong stabilizing effect *in cellulo*, suggesting that Vin is already of optimal stability. Only carbamylated Hpg-Vin showed reduced activation of T-cells, for reasons as yet unknown. The structural alteration resulting from citrullination, however, did show a drastic effect on T-cell activation: it was completely abolished. Whether this is due to citrullination within the DERAA epitope, remains to be elucidated.

**Figure 4:**
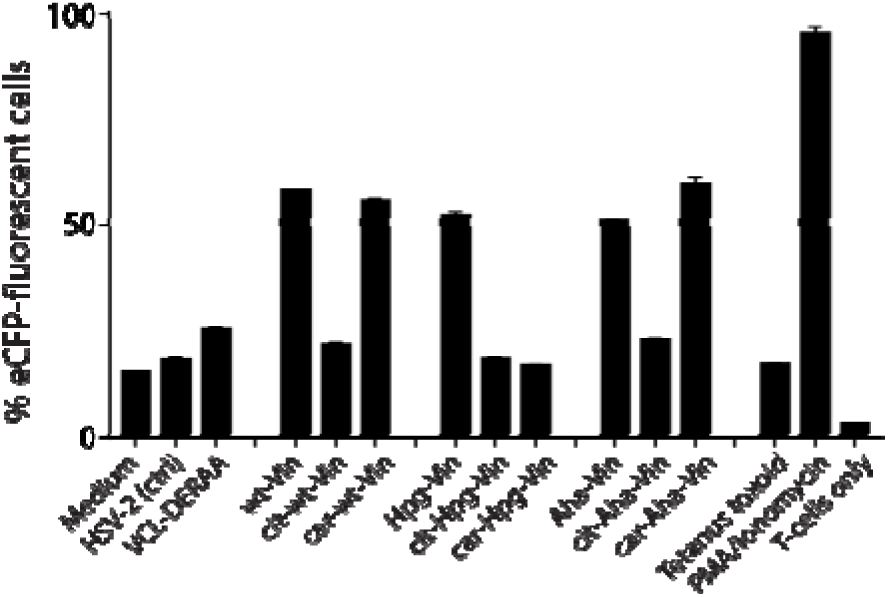
Activation of anti-DERAA T-cells by bioorthogonal vinculin derivatives. PBMC derived dendritic cells, isolated from HLA-DQ8+ donors were pulsed with peptides or Vin variants were cocultured with the Jurkat cells and used for stimulation. Activation of Vin-DERAA-specific Jurkat cells was measured with eCFP expression within the cells. The eCFP expression is dependent on NF_κ_B and is therefore a marker of TCR activation and costimulation, the percentage of live eCFP-positive cells is plotted in bars. HSV2 (Herpes Simplex Virus 2 peptide) and Tetanus toxoid protein are negative controls. VCL-DERAA peptide and PMA-ionomycin are the positive controls. Error bars show the SD. Data is representative of two experiments with biological triplicates.

## Conclusion

In summary, we were able to show that our bioorthogonal antigen variants are powerful tools to track the fate of antigens upon processing by APCs. They are structurally very similar to wild type antigens, the bioorthogonal groups are stable to the harsh conditions found in the lysosome, and the handles are not cleaved/destroyed by proteolysis. This means that – despite the added complication of an additional bioorthogonal ligation step - they are an excellent reagent set for the study of antigen processing and presentation. Also, no genetic manipulation of the immune cells is required, which we show make these reagents compatible with primary human immune cells.

Furthermore, the fact that nucleophilic sidechains remained available for post-translational modification now allows the study the effect of these modifications, such as carbamylation, but also acetylation, methylation, and non-enzymatic post-translational modifications, such as glycation on the rate of degradation/presentation in antigen presenting cells.

