## Supplementary Information for "Bioorthogonal antigens allow the unbiased study of antigen processing and presentation"

**Materials and Methods:**

**Chemicals**

Chemical reagents for buffer preparation and chemical synthesis were purchased from Acros (Belgium), Chem-Lab (Belgium), Honeywell Riedel-de Haën (Germany), Merck (The Netherlands), Novabiochem (The Netherlands), Sigma Aldrich (The Netherlands), Sigma Life Sciences (The Netherlands) or Sphaero Hispanagar (Spain) and used without further purification unless stated otherwise.

Fluorophores (Alexa Fluor 488 Azide, Alexa Fluor 488 Alkyne, Alexa Fluor 647 Azide and Alexa Fluor 647 Alkyne), were purchased from Thermo Fisher Scientific.

**Cell Culture**

**Bone Marrow Dendritic Cell Differentiation**

Bone marrow (BM) was isolated from 8-12 weeks old C57BL/6 mice kept under specific pathogen-free conditions (Strain: C57Bl/6NHsd, H-2^b^ Haplotype; Envigo Inc., Huntingdon, United Kingdom) essentially as described^1^. Femurs and tibiae of female or male mice were removed and surrounding tissue was removed. Thereafter, intact bones were left in 70% ethanol for 2–5 min and washed with sterile PBS. The bone marrow was flushed from femurs and tibia after removal of the bone-ends using pre-warmed IMDM (Sigma, ref# I3390) using a 20 mL Syringe with a 0.45 mm x 23 mm hypodermic needle (G26, Terumo Europe, NN-2623R). The BM was then homogenized through a 70 μm cell strainer (Falcon, ref# 352350).

The cells were then centrifuged 5 min at 350 *g* and resuspended in 10 mL IMDM supplemented with 8% heat-inactivated fetal calf serum (FCS, Sigma, ref# F0804, lot# 015M3344), 2 mM Glutamax^TM^ (GIBCO, ref# 35050-038), 20 μM β-Mercaptoethanol (Gibco, ref# 31350010), 50 IU/mL penicillin and 50 μg/mL streptomycin, and recombinant GM-CSF (20 ng/mL, Peprotech, ref# 315-03) to a concentration of 0.5 x 10^6^ cells/mL. The cells were incubated in non-adhesive petri dishes at 37°C, 5% CO_2_, under humidified air.

**OT-I and OT-II isolation and Culturing**

OT-I/CD45.1 mice, which have a transgenic Vα2Vβ5 TCR specific for the OVA_257–264_ epitope in association with H2-Kb , were bred in the specific pathogen-free animal facility of the Leiden University Medical Center (LUMC). The animal experiments have been reviewed and approved by the animal experimental committee of Leiden University OT-II mice (CD4^+^ T-cell transgenic mice expressing a TCR recognizing the OVA-derived T_h_ epitope ISQAVHAAHAEINEAGR in association with I-Ab)^2^ were bred and kept at the LUMC animal facility as well as at the Leiden Advanced Drug Research Centre (kindly provided by Dr. Bram Slutter) under specific pathogen-free conditions. All mice were used at 8–12 weeks of age.

Briefly, spleens from OT-I and OT-II TCR transgenic mice were harvested and homogenized using a 70 μm cell strainer. Untouched CD8^+^ or CD4^+^ T-cells were isolated with a Milteny mouse T-cells isolation kit for CD8^+^ or CD4^+^ T-cell negative selection, according to the manufacturer’s instructions. T-cell purity was typically higher than 85%.

**D1 dendritic cells**

D1 cells were cultured as described previously^3^. Briefly, cells were cultured in “complete IMDM” containing 10% heat-inactivated FCS, 100 IU/ml penicillin, 50 μg/ml streptomycin, 2 mM L-glutamine, and 50 μM β-mercaptoethanol. This medium was supplemented with 30% fibroblast supernatant from NIH/3T3 cells (collected and filtered from confluent cultures) containing 10-20 ng/ml mouse rGM-CSF. D1 culture conditions were 5% CO_2_ at 37°C.

**RAW 264.7 cells**

RAW macrophages were cultured as described previously^4^. Briefly, cells were cultured in DMEM (Sigma-Aldrich) containing 10% heat-inactivated FCS, 2 mM Glutamax^TM^, 50 IU/mL penicillin and 50 μg/mL streptomycin. RAW 264.7 culture conditions were 5% CO_2_ at 37°C.

**Human Cell Culture Protocols**

*Dendritic cell generation*

HLA-DRB1*04:01/-DQA1*03:01/-DQB1*03:02 (HLA-DR4/-DQ8)-positive buffy coats were received from Sanquin Bloodbank. Peripheral Blood Mononuclear Cells (PBMCs) were isolated with Ficoll-paque gradient and CD14^+^ cell isolation was performed with MACS beads according to the manufacturers protocol with slight alterations (Cat#: 130-097-052, Miltenyi Biotec). In short, PBMCs were incubated with 10 uL/10^7^ cells CD14^+^ microbeads and incubated for 15’ at 4°C. Coated PBMCs were run over an LS column twice to purify the CD14^+^ fraction. To differentiate the CD14^+^ cells to dendritic cells, the CD14^+^ cells were counted and seeded at 5x10^5^ cells/well in a 24-wells plate, in medium (IMDM, Cat#:12440-053, Gibco) supplemented with 1% Glutamax, 1% penicillin/streptomycin, 8% Fetal Calf Serum (FCS), 800 U/mL GM-CSF (Cat#:300-03, Peprotech) and 500 U/mL IL-4 (Cat#:200-04, Peprotech) and cultured for 6 days. After differentiation of the CD14^+^ cells to immature dendritic cells, the cells were fed with 200 μg POI per well and were allowed to take it up for ~18 hours. Peptides were added 1 hour before maturation at 10 µg/mL.

Subsequently, the dendritic cells were matured with a maturation cocktail: 100 ng/mL GM-CSF (Cat#: 300-03, Peprotech), 15 ng/mL TNFα (Cat#: 210-TA-005, R&D), 10 ng/mL IL-1β (Cat#: 201-LB-005, R&D), 10 ng/mL IL-6 (Cat#:200-06, Peprotech), 1 µg/mL PGE2 (Cat#: 14010, Cayman Chemical), 500 U/mL IFNγ (Cat#: 300-02, Peprotech). Dendritic cells were allowed to mature for ~30 hours before they were harvested and co-cultured with the T-cells.

*Jurkat Triple Parameter Reporter (TPR) cell line*

A Jurkat 76 cell line, positive for CD4 (by transduction) and NFAT-eGFP, NFkB-CFP and AP-1-mCherry (by transduction) was kindly provided by Mirjam Heemskerk from the LUMC.^5^^,^^6^ The cell line was transduced with Vin-DERAA-specific T-cell receptor in the group of Heemskerk by R. Hagedoorn, as has been previously described ^6^. The transduced Jurkat T-cells were selected on their TCR expression and sorted in single cells. One of these clones was shown to be highly responsive and was used for all future experiments described here. The JurkaT-cells are cultured in IMDM supplemented with 1% glutamax, 1% Penicillin/streptomycin and 8% FCS.

**Recombinant protein expression**

***Vinculin***

The plasmid (pET3d, Amp^R^, Novagen) encoding of wild type Vinculin_435-742_ (hereafter referred to as wt-Vin) was transformed into the methionine auxotroph *E.coli* B834(DE3) (met-aux, Genotype: F- ompT hsdSB (rB- mB-) gal dcm met(DE3), Novagen #ref 69041). The protein was expressed from the overnight culture of a single colony. 100 mL of this overnight culture (o.n., Ampicillin 50 µg/mL, 18 h, 37°C, and 150 rpm) was used for the inoculation per 1 L LB medium (Ampicillin 50 µg/mL, 37°C, 150 rpm). The cells were grown to an OD_600_ of 0.6-1.0 prior to the addition of 1 mM isopropyl-β-D-1-thiogalactopyranoside (IPTG) for 18 h (30°C, 150 rpm). The cells were harvested by centrifugation (4000 g, 20 min, 4°C) and washed once with Tris-buffered saline (TBS; 50 mM Tris-HCl, 300 mM NaCl, pH 8.0)

***Cell lysis and affinity purification of Vinculin_435-743_:***

Cell pellets were weighed, resuspended in TBS to 1-4 g/mL, prior to lysis by French Press (1.5-kbar, Stansted, Pressure Cell Homogeniser). Lysed cells were centrifuged (15,000 *g*, 1 h, 4°C). The soluble fraction was then loaded onto a Ni-NTA affinity resin (Thermo Fisher Scientific, catalogue no. 88221) pre-equilibrated with TBS. After incubation for 1 h under gentle rotation, the beads were first washed with 2 column volumes (CV) of washing buffer 1 (TBS, 10 mM Imidazole, pH 8.0) and subsequently with 3 CV washing buffer 2 (TBS with 50 mM Imidazole). The resin was then treated with elution buffer (TBS, 250 mM Imidazole, pH 8.0, 2 mL) to obtain the protein of interest (POI) in ~ 10 mg/mL, which was then exchanged into TBS using a Sephadex G25 resin (PD-10 column; GE Healthcare).

***Expression of bioorthogonal vinculin variants***

A single colony of *E. coli* B834 transformed with pET3d-Vin was grown in 50 mL LB augmented with 1% w/v glucose and grown overnight. The next morning, the culture was diluted 1:100 with fresh LB media and cells were grown to an OD_600_ of 0.5-0.6. The resulting culture was then centrifuged (2000 *g*, 10 min, 4°C) and the supernatant discarded. The cells were washed and resuspended in SelenoMet^TM^ media (Molecular Dimensions, USA) without additional methionine. The cells were incubated at 37°C for 30 min, followed by 30 minutes at 30°C (180 rpm) after which either _L_-Azidohomoalanine (Aha, 0.4 mM final concentration) or _D,L_-Homopropargylglycine (Hpg, 0.8 mM final concentration) was added. The cells were incubated for a further 30 minutes at 30°C (OD_600_ 0.7-1.0), the culture was induced with 1 mM IPTG for 18 h. Cell lysis and purification were then performed as described above.

Incorporation efficiency of the bioorthogonal amino acids was analyzed by LC-MS and in-gel fluorescence using bioorthogonal ligation with appropriately modified (i.e. Azide/Alkyne) Cy5 and Alexa 488 dyes.

***Ovalbumin expression***

The gene for the wt, full length Ovalbumin (hereafter referred to as wt-Ova, accession number V00383) was cloned into the pMCSG7 vector as described elsewhere^7^ and transformed into the methionine auxotroph expression strain, namely *E.coli* B834(DE3). The construct contained an N-terminal MHHHHHHSSGVDLGT***ENLYFG***SNA sequence for Ni-NTA purification (underlined) and a TEV-cleavage (italic bold) site. The expression was performed as described for wt-Vinculin.

***Cell Lysis and purification of Ova***

After the expression period, the cells were harvested by centrifugation (20 min, 3000 *g*). The pellet was resuspended in lysis buffer (50 mM NaH_2_PO_4_, 300 mM NaCl, 10 mM imidazole, pH 8.0, 10 % glycerol, 1 mg/mL Lysozyme (Amresco, ref#0663-5G), complete EDTA-free protease inhibitor (Roche, ref# 11836170001), β-mercaptoethanol (final concentration 5 mM), 250 U benzonase per 100 mL culture volume) to a final concentration of 5-10 g/mL. The cells were lysed with a French press at 1.5 kbar and the flow-through was centrifuged at 15000 g (30 min, 4°C) to separate soluble and insoluble components. The soluble-fraction was purified using nickel-affinity chromatography using a fast protein liquid chromatography (FPLC) system (Äkta Start Purification System, The Netherlands). The soluble-fraction was loaded onto a Ni-NTA Superflow cartage (Qiagen, ref # 30721) that was equilibrated with 10 CV buffer 1 (50 mM NaH_2_PO_4_, 300 mM NaCl, 10 mM imidazole, 10 % glycerol, pH 8.0). The column was washed with buffer 2 (10 CV, 50 mM NaH_2_PO_4_, 300 mM NaCl, 20 mM imidazole, 10 % glycerol, pH 8.0). Protein elution was then initiated by buffer 3 (50 mM NaH_2_PO_4_, 300 mM NaCl, 500 mM imidazole, 10 % glycerol, pH 8.0) using a 20 CV gradient (1 mL/min, 20 mL total volume), from 20 mM up to 250 mM imidazole (50% Buffer 3). All fractions were analyzed via SDS-PAGE and fractions containing >90% ovalbumin were collected and combined.

The protein was further purified by anion exchange chromatography. The combined fractions were dialyzed overnight into binding buffer (50 mM NaH_2_PO_4_, pH 8.0) and loaded onto an anion exchange column (GE Healthcare-HiTrap Q HP, ref# 17-1153-01) that had been equilibrated with 10 CV binding buffer. The protein was eluted using an elution buffer gradient (50 mM NaH_2_PO_4_, 0-500 mM NaCl, pH 8.0). Elution fractions were analyzed by SDS-PAGE and the fractions containing the POI were combined and dialyzed o.n. into 10-100 mM NaHCO_3_, pH 8.0 to yield 2 mL of a 2-5 mg/mL solution of wt-Ova (yield 0.4-2 mg/L culture). The protein solution was snap-frozen and stored at -80°C until further use.

***Expression of bioorthogonal Ova variants***

Bioorthogonal protein expression was performed as described for the wt-Ova protein with the following changes. After reaching the optimal optical density of 0.6-0.9 for the pre-culture, the medium was exchanged for the SelenoMet methionine deficient medium and the culture was left to grow for another 30 min at 37°C. Then the medium was supplemented with unnatural amino acid, namely, Aha (0.4 mM) or D-/L-Hpg (0.8 mM) and expression was induced by addition of 1 mM IPTG. Expression was continued o.n. (30°C, 180 rpm). The culture was harvested by centrifugation (30 min, 4 °C and 4000 g). The cells were lysed as above and purified as described for wt-Ova.

**Enzymatic and Chemical Modification of Recombinant Proteins**

***In vitro* Carbamylation of recombinant proteins**

Vin variants were carbamylated using a protocol adapted from Fando *et al.*^8^. Briefly, the POI (2 mg/mL) was reacted with 0.2 M potassium cyanate (KOCN) in H_2_O in a 1:1 v/v ratio for 24 h at 37°C. Typical reaction volumes were 0.5-1.0 mL. The reaction progress was analyzed by LC-MS (Waters UPLC). Upon completion of the reaction, carbamylated proteins were purified by gel filtration over Sephadex G-25 (PD-10 prepacked column, GE Healthcare). Purified proteins were stored at -80°C until further use.

***In vitro* Citrullination of Recombinant Proteins**

Recombinant Vin variants were citrullinated as follows; 1 mg/mL of POI was dissolved in Tris-HCl (100 mM, pH 7.6) buffer containing CaCl_2_ in ddH_2_O (10 mM) and DTT (5 mM). Recombinant PAD4 (Sigma Aldrich, final concentration 1U) was added to the reaction mixture. Samples were incubated at 37°C for 5 h-o.n. under constant shaking (400 rpm). Upon completion of the reaction, citrullinated protein samples were purified via gel filtration and stored at -80°C for further use.

**DEGRADATION ASSAYS**

**Protein degradation in vitro using lysosomes**

For magnetic lysosomal extraction, the protocol described by Walker and Lloyd-Evans with several modifications was used.^9^ Briefly, 20*10^6^ RAW macrophages were pulsed for 24 h at 37°C with 10% dextran coated magnetite (DexoMAG 40, Liquids Research Ltd) nanoparticles. After rinsing with PBS, cells were incubated with complete DMEM for another 24 h. At the end of this chase period, the cells were rinsed and harvested in PBS, centrifuged (200 *g*, 5 min), and homogenized with buffer 1 (15 mM KCl, 1.5 mM MgCl_2_, 10 mM HEPES) using a dounce homogenizer with 30 cycles. Following homogenization, buffer 2 ( 200 mM HEPES pH 7.2, 0.1 mM sucrose, 375 mM KCl, 22,5 mM MgCl_2_ and 25 µL Benzonase in 5 mL) was added to the cells. Subsequently, cells were centrifuged at 200 *g* for 10 min and the supernatant was loaded on a buffer 1 equilibrated LS column (Miltenyi Biotec, Auburn, CA) placed inside a strong magnetic field (QuadroMACS Separator). Note that following steps were conducted at 4 ℃. The non-magnetic fraction was collected through washing the column with cold buffer 1 (2x 5 mL). Next the LS column was washed with 1 mL of PBS with 0.1 mM sucrose and Benzonase (25 U) to detach nuclear fractions. Thereafter, the LS column was washed with 1 mL of PBS containing0.1 mM sucrose and separated from the magnetic field to elute the magnetically labeled lysosomes with 0.5-1 mL of PBS and 0.1 mM sucrose buffer. For long term storage, the lysosomes are snap-frozen in liquid N_2_ and could be stored up to 3 months at -80 ℃ without loss of proteolytical activity.

In a typical degradation experiment, lysosomes were subjected to multiple rapid (subsequent 4-5 times) freeze and thaw cycles to lyse them. Prior to use, the protein concentration was determined using the Bradford or Qubit protein assay. Approximately 18 µg of lysosomal proteins were used for each reaction condition. Recombinant or purchased proteins were diluted to a final concentration of 15 µM in a total reaction volume of 200 µL. Reaction mixtures were acidified with 0.4 µL 5 M HCl to pH 5.0 and samples were taken after 0, 15, 30 min as well as 1, 2, 4, 18, 22 and 48 h. Collected samples were analyzed via SDS-PAGE and the specific protein amount was visualized by Coomassie staining.

**Bioorthogonal protein degradation *in cellulo***

Differentiated BMDCs or immature D1 cells were seeded in a tissue culture treated 24-well microtiter plate at 0.2 x 10^6^ cells/well in 250 µL medium/well and were left to attach for 3 h. The bioorthogonal antigens dissolved in IMDM complete medium were added at the indicated concentrations over 1 h (pulse, BMDCs) or 2 h (pulse, D1 cells) at 37°C, 5% CO_2_ under humidified air. Then the medium was aspirated gently, and the cells washed with pre-warmed IMDM medium. Subsequently, cells were pulsed for the indicated times (varying from 0-48 h). The supernatant was then aspirated, and the cells resuspended in 100 μL of lysis buffer (100 mM Tris-HCl pH 7.5, 50 or 150 mM NaCl, complete protease inhibitor cocktail (EDTA-free), 0.25% CHAPS, 250 U Benzonase (Sigma, ref# E1014-25KU)) and incubated for 30 mins. The resulting samples were normalized by total protein concentration. For this purpose, the lysate was cleared via centrifugation (20000 g, 1 h) and subjected either to Bradford assay (Biorad, the Netherlands, ref# 5000001) or Qubit Protein Assay (Thermo Fisher, the Netherlands, ref# Q33211)

20 μL of the normalized lysate was then mixed with 20 μL CuAAC-buffer. This buffer was generated via addition of following chemicals in the particular order: CuSO_4_ in Milli-Q water (100 mM stock concentration, 6.4 mM final concentration), sodium ascorbate in Milli-Q water (1 M stock concentration, 37.5 mM final concentration), followed by 2,4,6-Trimethoxybenzylidene-1,1,1-tris(hydroxymethyl) ethane methacrylate (TTMA) from a DMSO stock (100 mM, 10 μL, 1.3 mM final concentration) and HEPES (stock concentration 100 mM, final concentration 88 mM, pH 8.0). Finally, fluorophore-alkyne or –azides were added from DMSO-stocks (stock concentration 2 mM, 1-2 μL, 2.5-5 µM final concentration). Please note, upon addition of sodium ascorbate to CuSO_4_, a color change from blue to brown (reduction of copper(II) into copper(I)) and finally to yellow should occur. Samples were incubated for 30 minutes at r.t. under gentile agitation (100-200 rpm) in the dark prior to the addition of Laemmli-buffer and SDS-PAGE analysis after which in-gel fluorescence was measured at the respective wavelengths and the specific protein amount was visualized by Coomassie staining.

**T-CELL PROLIFERATION ASSAYS**

***In vitro* MHC II-restricted Antigen Presentation Assay**

BMDCs were seeded (50.000 cells/well, medium described above) in treated cell culture 96-well plates (flat bottom) and left at 37°C for 2 h to allow adhesion. Then, cells were incubated with different concentrations of wild type and bioorthogonal versions, ranging from 0.4 to 93.3 µg/mL, for 6 h. Stock solutions of proteins were prepared in 10 mM NaHCO_3_ and DMEM complete medium. The class II epitope of OVA (T_h_ epitope ISQAVHAAHAEINEAGR), was used as a positive control for antigen presentation. After the period of incubation, cells were washed with warm IMDM and CD4^+^ OTII cells were added (50,000 cells/well, RPMI medium supplemented with 10% heat inactivated FCS, 2 mM glutamax, 50 μg/ml penicillin/streptomycin, 20 μM of β-mercaptoethanol). As a read out for T-cell activation, IL-2 secretion (ELISA) was measured in the supernatants 20 h later. Assays were performed in duplicates.

**Human T-cell activation assay (Dendritic cell and Jurkat T-cell co-culture and evaluation of Jurkat activation)**

Harvested dendritic cells were transferred to a 96-wells plate, 3 wells per condition, except for the Jurkat only conditions. When the HLA-DQ/TCR interaction was blocked with antibodies, the dendritic cells were pre-incubated for 1 hour with 20 µg/mL anti-HLA-DQ (SPVL3) and anti-HLA-DR (B8.11.2) antibodies before Jurkat T-cells were added. Upon addition of Jurkat T-cells to the wells in a 1:1 or a 1:2 ratio (even number of T-cells), the cells were cultured together for ~16-18 hours before the cells were harvested, stained with Zombie NIR Fixable viability dye (Cat#: 423105, BioLegend) according to manufacturer’s protocol and fixed with 3% PFA. Cells were measured using the BD LSRFortessa. Results were analyzed using FlowJo Version 10.4.2.

**CONFOCAL MICROSCOPY**

Differentiated D1s were seeded in a tissue culture treated, poly-lysine coated 96-well microtiter plate at 5 x 10^4^ cells/well and were left to attach 3 h. Bioorthogonal antigens were diluted to a final concentration of 2 mg/mL, of which 50 µg was added to each well bearing 200 µL of cell suspension over 2 h (pulse) and were chased for 0, 2, 4 and 24 h at 37°C, 5% CO_2_ under humidified air. Cells were washed with PBS before fixing with cold 2% PFA in PBS for 30 minutes at room temperature (r.t.) or with 0.5% PFA in PBS o.n.. After fixation, cells were again washed with PBS, then with the quenching buffer (20 mM glycine in PBS) and permeabilized with staining buffer (0.1% IGEPAL, 1% BSA in PBS) for 20 minutes at room temperature. After washing with PBS and subsequently with 100 mM HEPES (pH 8), bioorthogonal antigens were incubated 1 h with click mix (1 mM CuSO_4_, 10 mM sodium ascorbate, 1 mM THPTA ligand, 10 mM aminoguanidine, 100 mM HEPES pH 8 and 5 μM Alexa Fluor 647-azide or alkyne. Upon CuAAC, cells were washed first with 100 mM HEPES (pH 8), then with PBS, then incubated for 30 minutes with 1% BSA in PBS and stained with Alexa Fluor ® 488 LAMP-1 antibody (Biolegend, clone 1D4B, Cat#:121608, 1:200 dilution, staining in 1% BSA, 0.1% tween-20 in PBS) for 1h. Cells were washed three times with PBS before addition of fluoroshield mounting medium containing DAPI (Cat#: ab104139) and samples were imaged with a Leica SP (63x oil lens, N.A.=1.4) or Andor Dragonfly confocal microscope.

**ANALYTICAL METHODS**

***SDS-PAGE analysis*** ^10^

For SDS-PAGE analysis all samples were heated for 5 minutes at 95°C (exception: samples containing click cocktail). 20 μL of each sample was loaded onto a 15% SDS-PAGE gel (0.75 or 1.5 mm) and run for ~70 min at constant 170 V. Subsequently in-gel fluorescence was measured at indicated wavelength filters for Alexa 488, Alexa 647 or Cy5, after which a Coomassie staining was performed. For imaging the gels, Biorad Chemidoc Imager and ImageLab 5.2 software (Biorad) was used.

***NATIVE PAGE analysis***

Native PAGE was performed as described by Arndt *et al.*^11^. Images were taken via Chemidoc Imager and were analyzed via ImageLab 5.2 software (Biorad).

***Circular Dichroism (CD)***

All recombinant wt and bioorthogonal Vin and Ova variants were characterized via CD spectroscopy. Far UV-CD spectra were recorded using a Jasco J815 CD spectrometer equipped with a Jasco PTC 123 Peltier temperature controller (Easton, MD) between 190-260 nm. A minimum of five spectra with an acquisition time of 70 seconds (s) for each scan in a 1 mm quartz cuvette at 1 nm resolution were acquired at r.t. and averaged. Typical protein concentrations were between 0.1-0.3 mg/mL.

Organic Synthesis

NMR spectra were measured on a Bruker AV-400MHz spectrometer at ambient temperature at the Leiden Institute of Chemistry NMR Facility. Chemical shifts were recorded in ppm. Residual solvent peaks were used as an internal standard.

*Synthesis of L-Azidohomoalanine (_L_-AHA)*

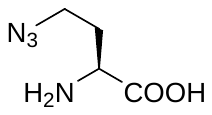
L-Aha was synthesized according to a previously described procedure ^12^. ^1^H-NMR (D_2_O), 400 MHz: δ [ppm] = 4.05 (t, 1H, α-CH), 3.55 (t, 2H, γ-CH_2_), 2.15 (m, 2H, β-CH_2_).

*Synthesis of D,L-Homopropargylglycine (_D,L_-Hpg)*

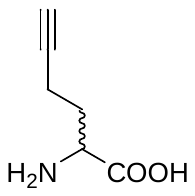

D,L-Hpg was synthesized according to previously described procedure ^13^. ^1^H-NMR (D_2_O), 400 MHz: δ [ppm] = 4.12 (t, 1H, α-CH), 2.3-2.12 (m, 1H), 2.11-2.0 (m, 1H).

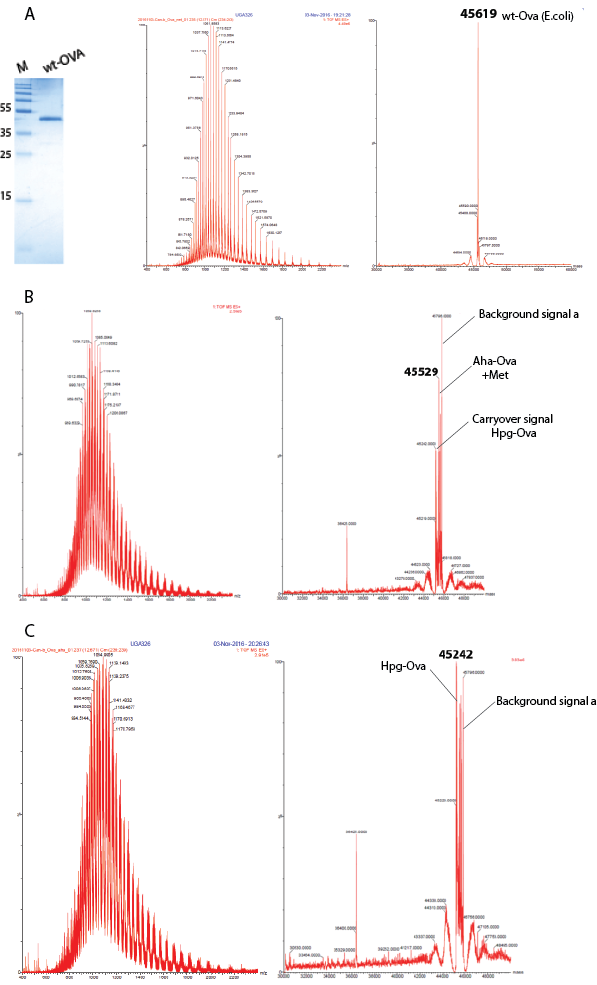

Figure S1: *Characterization of wt and bioorthogonal Ova derivatives*. A: left panel; SDS-PAGE analysis of wt-Ova (lane 2), right panel; TOF-MS analysis of wt-OVA (expected mass: 45618.7 Da, observed mass: 45619 Da), B: TOF-MS analysis of Aha-Ova (expected mass: 45534, observed mass: 45529 Da), C: TOF-MS analysis of Hpg-OVA (expected mass:45245 Da, observed mass: 45242 Da). Background signal refers to internal background noise due to low ionization energy.

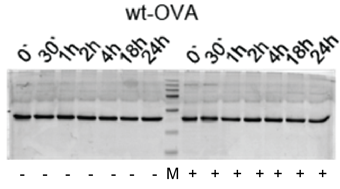

Figure S2: *Proteolysis of wt-OVA in vitro* wt-OVA was incubated with (+)/without(-) lysosomes extracts for 0 min, 30 min, 1 h, 2 h, 4 h, 18 h and 24 h. a total amount of 2.5 µg protein was loaded. No degradation was observed at any time point.

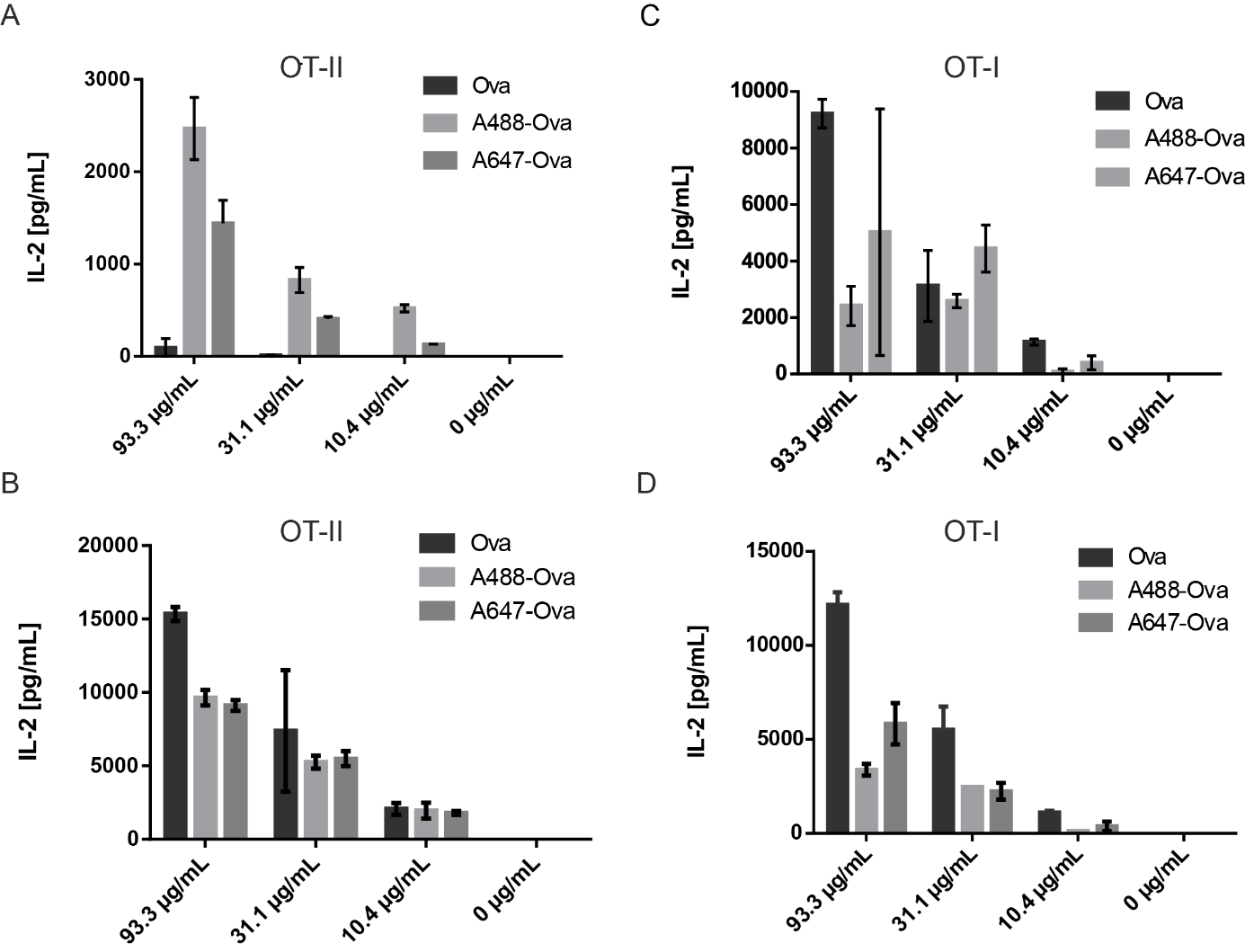

Figure S3: *Influence of fluorophore modification of Ovalbumin on the activation of antigen specific CD4+(OT-II) and CD8+ (OT-I) T-cells.* A-B: ELISA (IL-2) read out for CD4^+^ T-cell proliferation after feeding BMDCs with glycosylated commercially available Ova (Ova, dark grey) and fluorophore modified Ova (light grey, A488-Ova; grey, A647-Ova), n=1 with three technical replicates. C-D: ELISA (IL-2) read out for CD8^+^ T-cell activation after feeding BMDCs with glycosylated commercially available Ova (Ova, dark grey) and fluorophore modified Ova (light grey, A488-Ova; grey, A647-Ova), n=3 independent with at last two experimental (N=2) replicates for B-D.

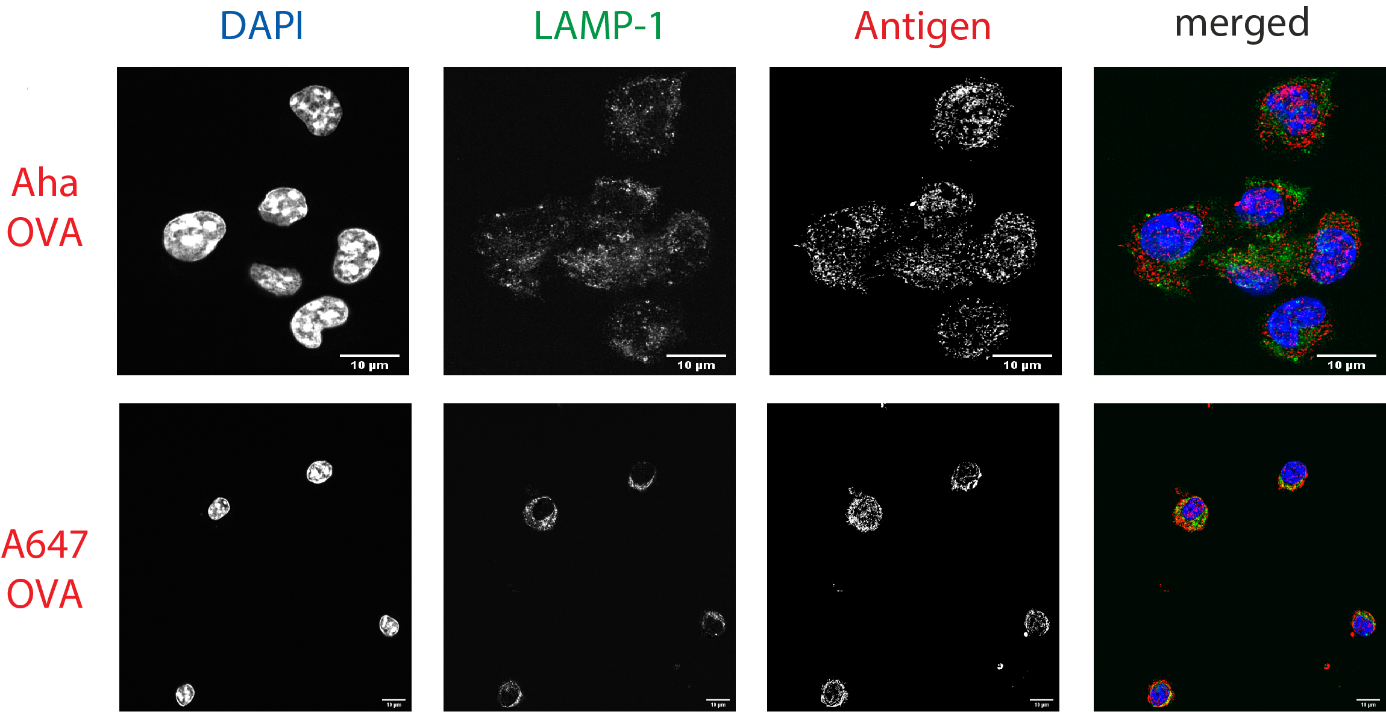

**Figure S4**: ***Imaging processing of bioorthogonal and fluorophore-modified Ova in D1-APCs via confocal microscopy*.** DCs were incubated for 2 h (pulse) with Aha or A647-Ova. Cells were fixed with 0.5% PFA and processed for immunofluorescence with the following primary antibodies. The nucleus was stained with DAPI (blue in merged images) and LAMP-1 (green in merged images) was used as a lysosomal marker. Aha-Ova was stained using copper catalyzed Huisgen cycloaddition (ccHc) with Alexa647-alkyne fluorophore (red in merged images). Upper panel: imaging of Aha-Ova, lower panel: imaging of A647-Ova. Scale bar is 10 µm (white bar, right corner).

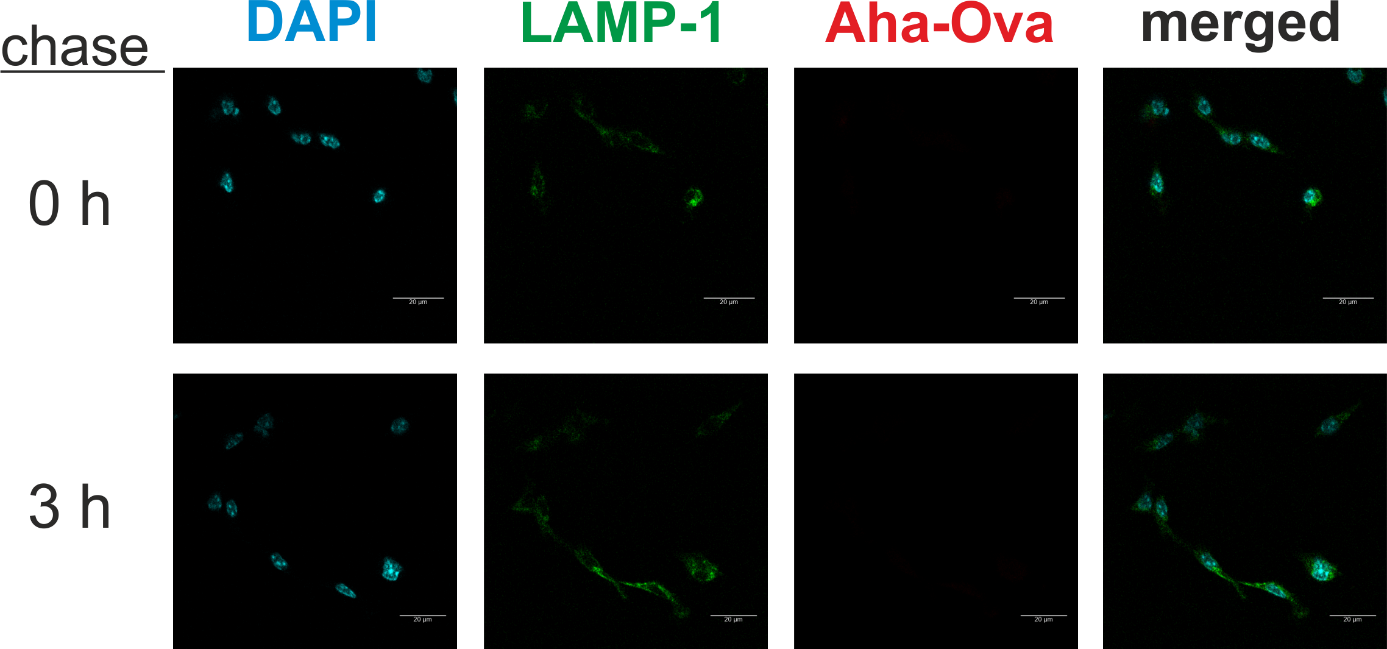

**Figure S5: *Control experiment for processing of wt-Ova in D1-APCs via confocal microscopy*** DCs were incubated for 2 h (pulse) with wt-Ova. Cells were chased for distinct timepoints (0 and 3 h), fixed with 2% PFA and processed for immunofluorescence. The nucleus was stained with DAPI (cyan) and anti-LAMP-1 (green) was used as a lysosomal marker. Wt-Ova was stained using ccHc with Alexa647-alkyne fluorophore (Thermo Fisher). Upper panel: Antigen uptake and processing after 0 h, lower panel: 3 h. The scale bar is 20 µm (white bar, right corner).

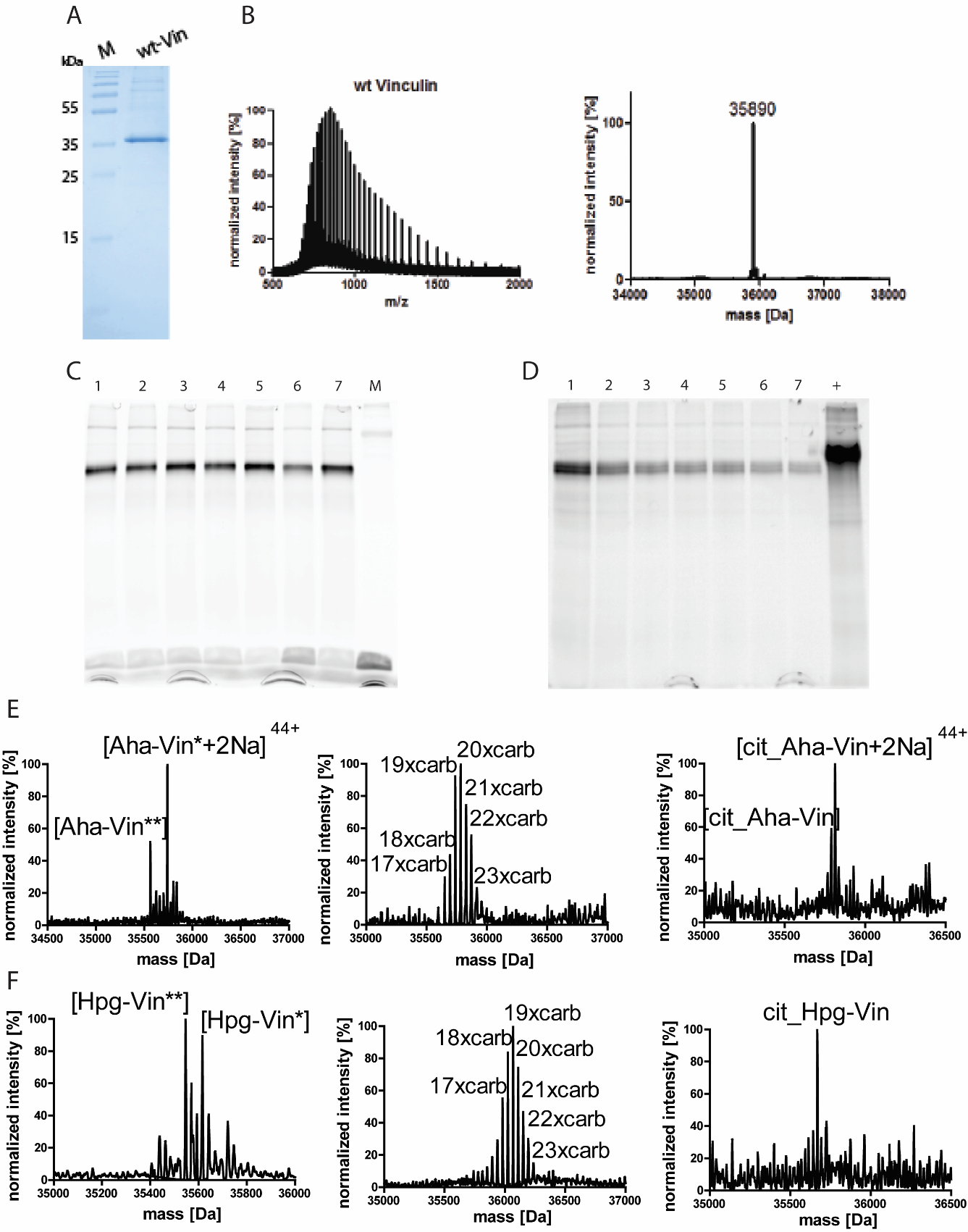

Figure S6: *Characterization of wt Vin & modified bioorthogonal Vin derivatives* A: SDS-PAGE analysis of wt Vin (lane 2), B) TOF-MS analysis of wt-Vin (expected mass: 35886 Da, observed mass: 35890 Da), C: In-gel fluorescence analysis of Aha incorporation at different ratios of fluorophore to protein concentrations, lane 1: fluorophore 1 eq: protein 1 eq., lane 2: fluorophore 1eq. : protein 3eq., lane 3: fluorophore 1eq. : protein 4eq., lane 4: fluorophore 1eq. : protein 5eq., lane 5: fluorophore 1eq. : protein 6eq., lane 6: fluorophore 1eq. : protein 7eq., lane 7: fluorophore 1eq. : protein 8eq., M: protein ladder. D: In-gel fluorescence analysis of Hpg incorporation at different ratios of fluorophore to protein concentrations, lane 1: fluorophore 1 eq.: protein 1 eq., lane 2: fluorophore 1eq. : protein 3eq., lane 3: fluorophore 1eq. : protein 4eq., lane 4: fluorophore 1eq. : protein 5eq., lane 5: fluorophore 1eq. : protein 6eq., lane 6: fluorophore 1eq. : protein 7eq., lane 7: fluorophore 1eq. : protein 8eq., +: positive control (Ova-azide). E: TOF-MS analysis of biorthogonal Aha-Vin (left, expected mass: 35715 Da, observed mass: 35714.6 for Aha-Vin** & 35759 Da for [Aha-Vin*+2Na]^44+^), and Hpg-Vin (right, expected mass: 35579 Da, observed mass: 35578.6 for Hpg-Vin** & 35759 Da for [Hpg-Vin*+2Na]^44+^). F: TOF-MS analysis of citrullinated bioorthogonal Vinculin derivatives, left; citrullinated Aha-Vin (expected mass: 35734 Da, observed mass: 35736 Da), right; citrullinated Hpg-Vin (expected mass: 35669 Da, observed mass: 35668 Da).

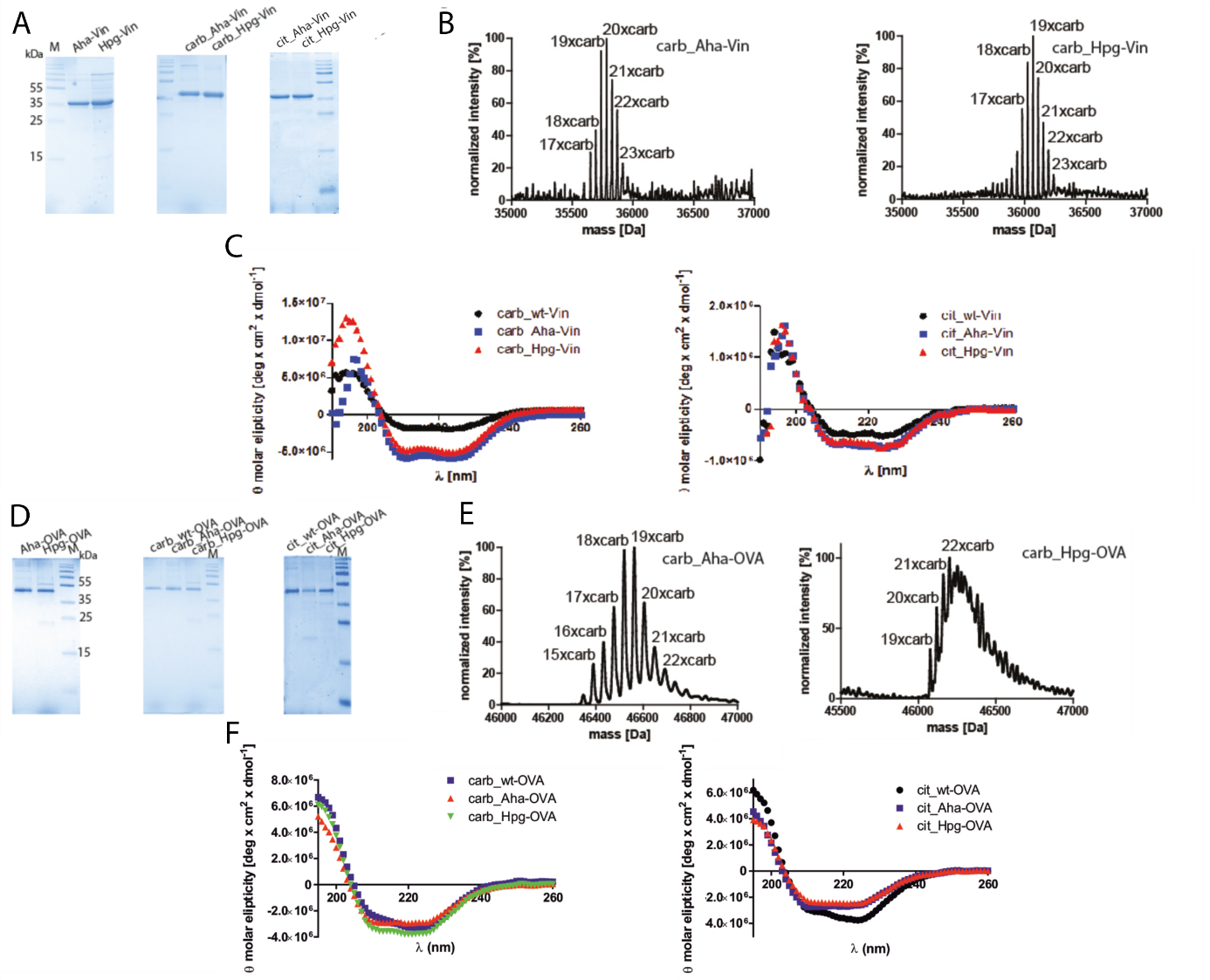

Figure S7: *Carbamylation and citrullination of (bioorthogonal) Vinculin.* A: SDS-PAGE analysis of purified, non-modified (left), carbamylated (middle) or citrullinated (right) bioorthogonal Vin derivatives. B: TOF-MS analysis of carbamylated bioorthogonal Vin derivatives, left; carb_Aha-Vin (mixed species from 17x-to 23x-carbamylated Aha-Vin, no traces of non-carbamylated Aha-Vin), right; carb_Hpg-Vin (mixed species from 17x-to 23x-carbamylated Hpg-Vin, no traces of non-carbamylated Hpg-Vin) C: Biophysical characterization of modified bioorthogonal Vin via circular dichroism (CD) spectroscopy, left panel: carbamylated Vin derivatives, right panel: citrullinated Vin derivatives. wt- (black circles), Aha- (blue squares) and Hpg-Vin (red triangles)

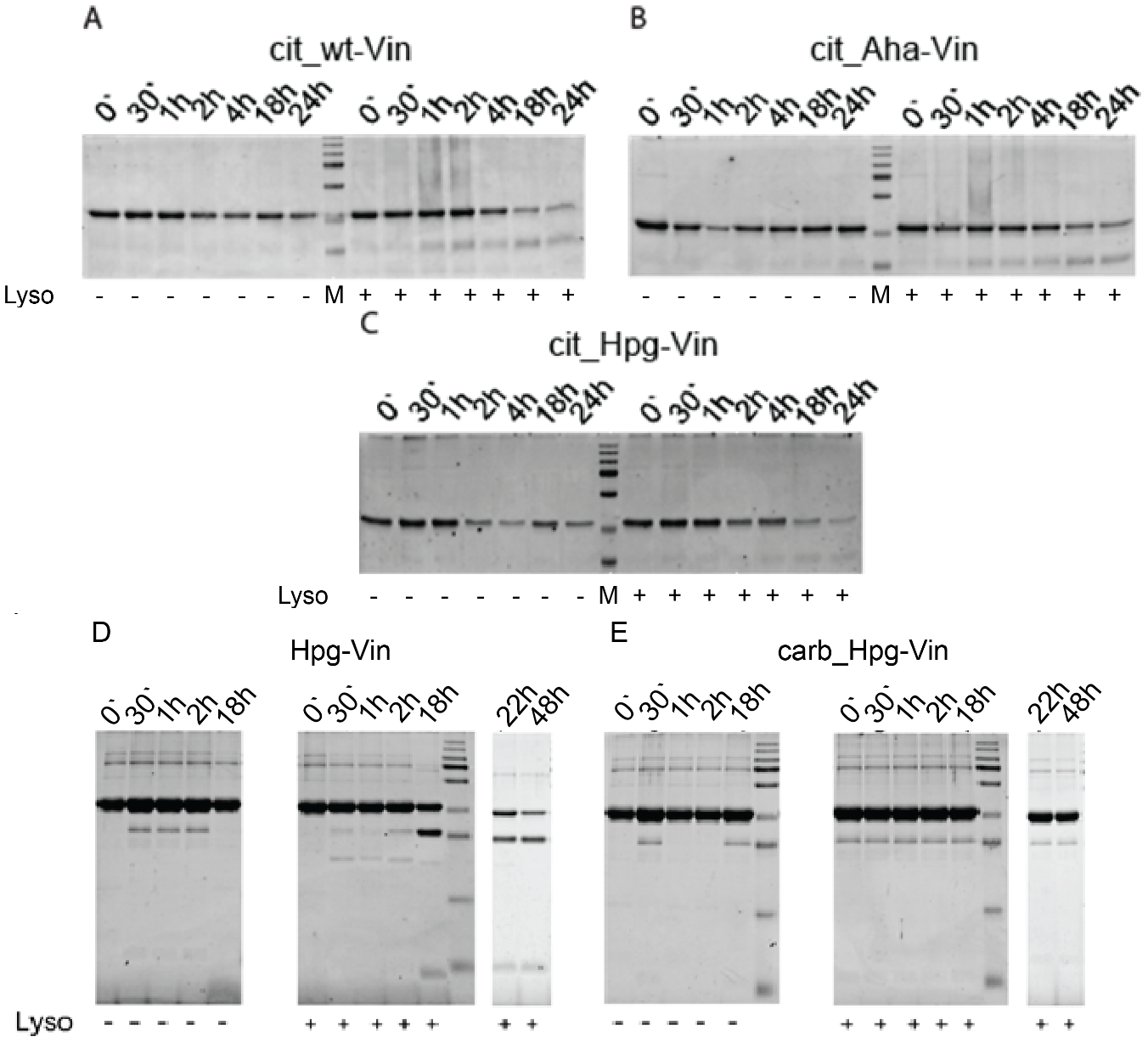

Figure S8: *Proteolysis citrullinated Vin, carb-Hpg-Vin and wt-OVA in vitro* A) cit_wt-Vin incubated with lysosomes extracts (Lyso +) for 0h, 30min, 1h, 2h, 3h, 18h and 24h. B) cit_Aha-Vin incubated with lysosomes extracts for 0h, 30min, 1h, 2h, 3h, 18h and 24h. (Lyso -) : samples not incubated with lysosomes (control samples) C) cit_Hpg-Vin incubated with lysosomes extracts for 0h, 30min, 1h, 2h, 4h, 18h and 24h. D) carb_Aha-Vin incubated with lysosomes extracts for 0h, 30min, 1h, 2h, 4h, 18h and 24h. D) Hpg-Vin incubated with lysosomes extracts for 0h, 30 min, 1h, 2h and 18h. Left; Hpg-Vin without lysosomes (control samples), right; Hpg-Vin incubated with lysosomes. E) carb_Hpg-Vin incubated with lysosomes extracts for 0h, 30 min, 1h, 2h and 18h. Left; carb_Hpg-Vin without lysosomes (control samples), right; carb_Hpg-Vin incubated with lysosomes. For all lanes on A-C a total amount of 2.5 µg protein was loaded. Total amount of protein/lane on D-E is 5 µg.

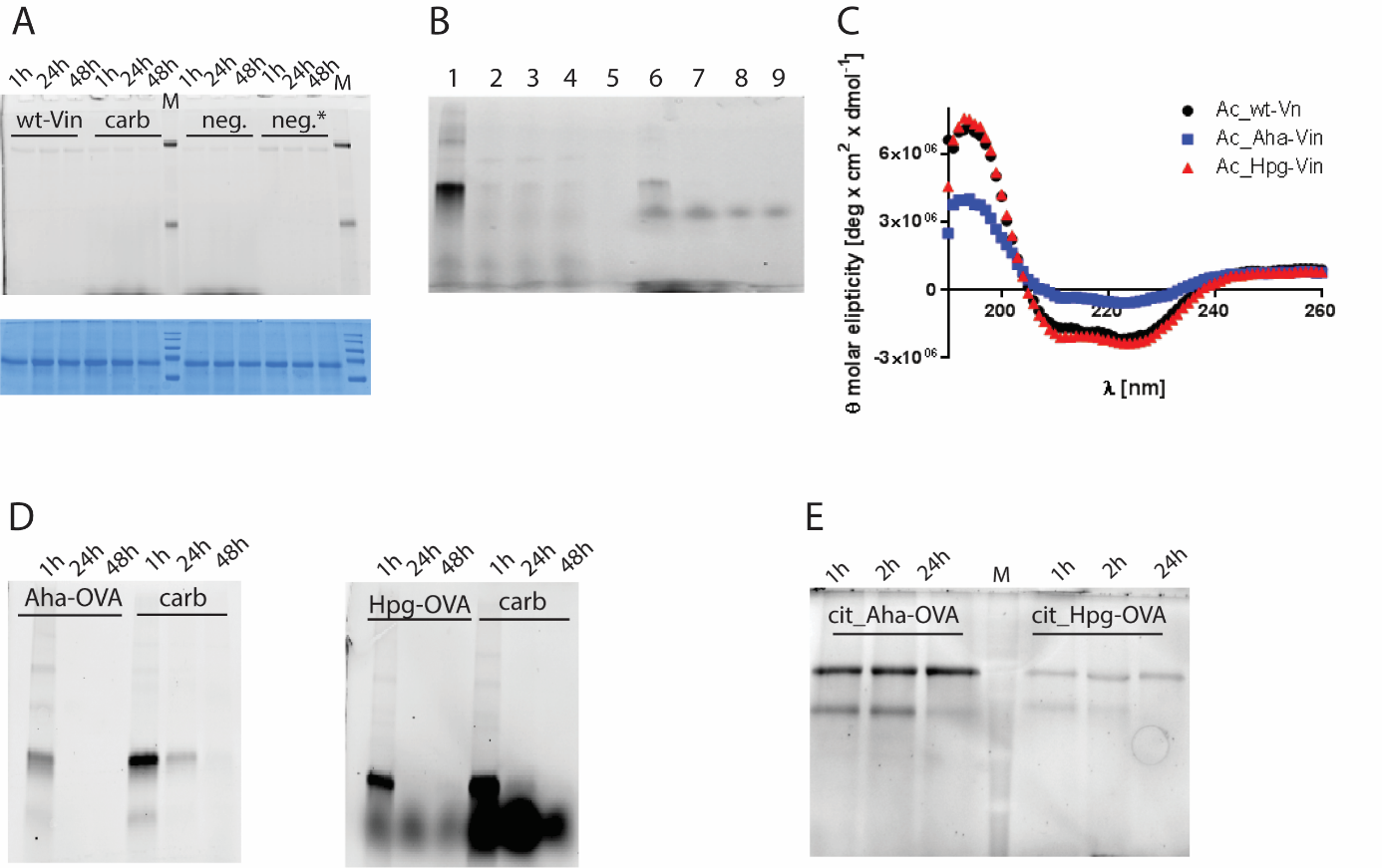

Figure S9: *In cellulo degradation of wt-Vin after uptake in BMDCs*. Cells were pulsed with wild type and carbamylated antigens for two hours and chased for the indicated time points. Subsequently the cells were collected, lysed and subjected to ccHc conditions to introduce AlexaFluor-488 with azide (neg.) or alkyne (neg.*) functionality. A: Degradation of wt (left) and carbamylated wt-Vin (grey: fluorescent signal from AlexaFluor-488; blue: Coomassie loading control). As expected, no fluorescent signal was observed.

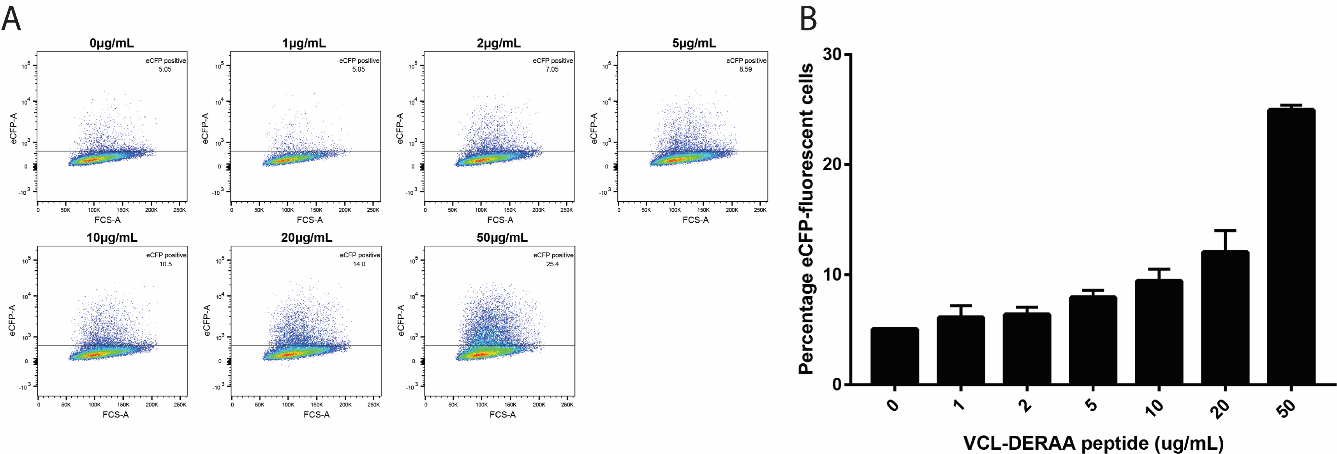

Figure S10: *T-cell activation of Jurkat-DERAA cells with titrated amounts of VCL-DERAA peptide.*

The T-cells were stimulated by HLA-DQA1*03:01/DQB1*03:02+ (HLA-DQ8+) EBV-transformed B cells presenting VCL-DERAA peptide with titrated amounts (0-50 µg/mL). The TCR on Jurkat-DERAA recognizes VCL-DERAA peptide in a dose-dependent manner. A: eCFP positivity in FACS was used as readout, B: Plotted data in bar graphs, data is shown as n=2 with biological triplicates. Error bars are presented as ±SD.

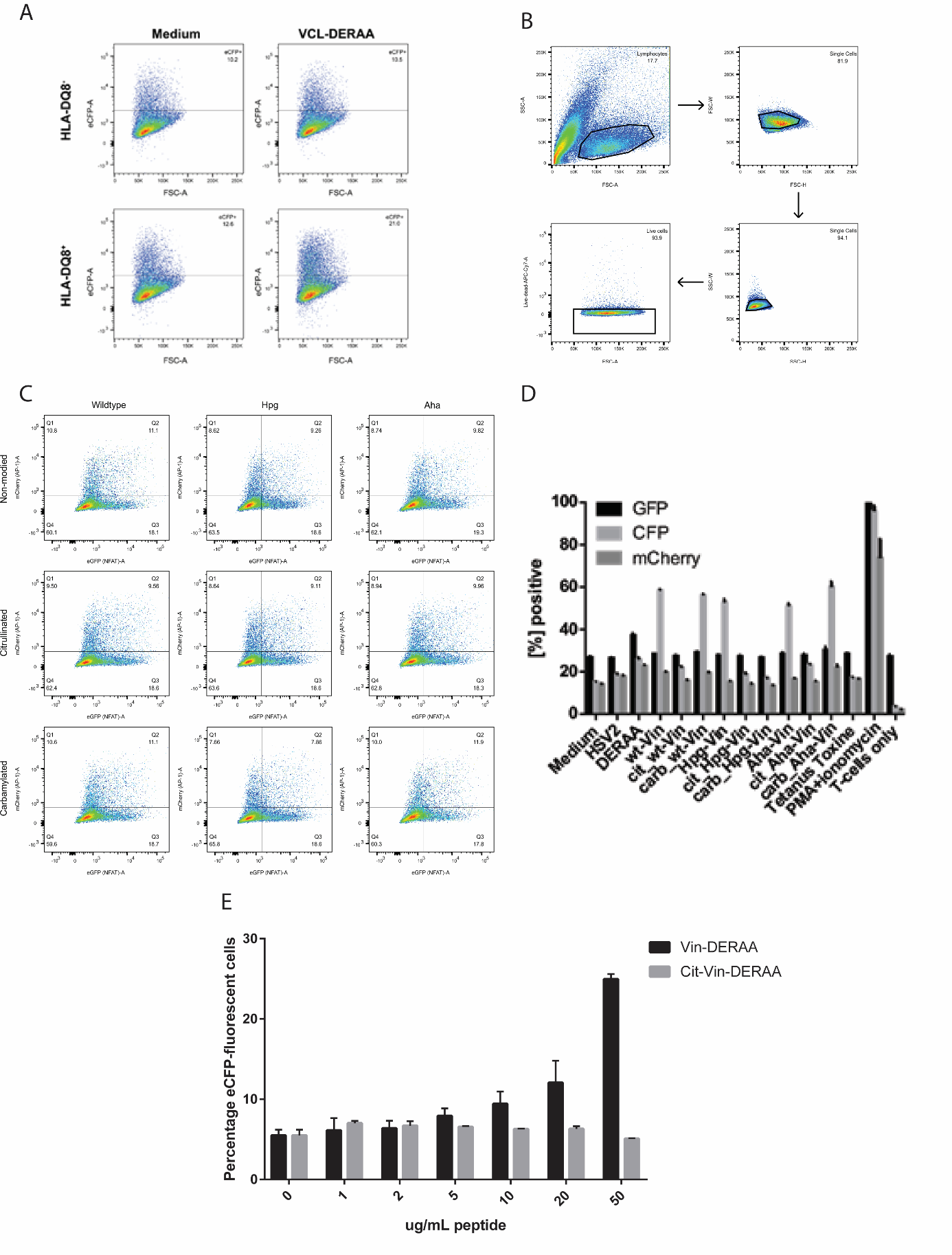

Figure S11: *FACS read-out for T-cell activation with VCL-DERAA peptide and bioorthogonal Vin derivatives.* A: FACS read-out (eCFP) for incubation of Jurkat-DERAA cells with VCL-DERAA peptide fed to HLA-DQ8^-^ B-cells (upper panel) and HLA-DQ8^+^ B-cells (lower panel), B: General gating strategy for Jurkat T-cell line (J76) with a focus on the single and living cells in the Jurkat T-cell population, C: T-cell activation assay with bioorthogonal antigens (eGFP expression as a readout). D: Double positivity in fluorescence for mCherry and CFP as a measure of T-cell response, plotted as bar graphs. Error bars represent ±SD. E: Titration experiments with Vin-DERAA and citrullinated Vin-DERAA (Cit-Vin-DERAA) peptide, percentage of eCFP positive cells was read out as a measure for T-cell activation.
